# Simulations reveal how touchdown kinematic variables affect top sprinting speed: implications for coaching

**DOI:** 10.1101/2024.10.08.617292

**Authors:** Nicos Haralabidis, Ashton J. Eaton, Scott L. Delp, Jennifer L. Hicks

**Author notes:** **Corresponding Author** Scott L. Delp, Stanford University, Stanford, CA 94305, USA.

## Abstract

Sprint performance is a priority for coaches and athletes. Several kinematic variables, including horizontal touchdown distance (HTD) and inter-knee touchdown distance (IKTD), are targeted by coaches to increase top sprinting speed. However, the results of past research are conflicting, potentially due to the use of experimental inter-athlete study designs where it is not possible to establish cause-effect relationships. In this study, we used a predictive simulation approach to assess cause-effect relationships between HTD and IKTD and sprinting speed. We scaled a three-dimensional musculoskeletal model to match the anthropometry of an international caliber male sprinter, and generated predictive simulations of a single symmetric step of top-speed sprinting using a direct collocation optimal control framework. We first used our simulation framework to establish the model’s top speed with minimal constraints on touchdown kinematics (the optimal simulation). Then, in additional simulations we enforced specific HTD or IKTD values (± 2, 4 and 6 cm compared to optimal). The model achieved a top speed of 11.85 m/s in the optimal simulation. Shortening HTD by 6 cm reduced speed by 7.3%, while lengthening HTD by 6 cm had a smaller impact on speed, with a 1.6% reduction. Speed in the simulation was insensitive to the IKTD changes we tested. The results of our simulations indicate there is an optimal HTD to maximize sprinting speed, providing support for coaches and athletes to adjust this technique variable. Conversely, our results do not provide evidence to support utilizing IKTD as a key technique variable for speed enhancement. We share the simulation framework so researchers can explore the effects of additional modifications on sprinting performance (https://github.com/nicos1993/Pred_Sim_Sprinting).

## Introduction

Athletes sprint to move a short distance at high speed, and the outcomes of numerous sporting scenarios can be affected by an athlete’s sprint performance. Sprint performance depends on the ability to rapidly accelerate and reach top speed. Several studies (1–3) have demonstrated the importance of top sprinting speed for the short distance track events, with top speed associated with faster 100 m times. Furthermore, in team sports, such as American football, soccer, and rugby, a greater top sprinting speed can affect the outcome of decisive attacking and defensive actions (4, 5). Thus, for coaches and athletes, increasing top sprinting speed is a sought-after goal.

While a large number of kinematic variables have been studied to understand the biomechanics of top-speed sprinting, coaches typically rely on a smaller subset of variables (6–8). One popular variable is horizontal touchdown distance (HTD) (Figure 1A): it is recommended that the plant foot should be only slightly ahead of the whole-body center of mass (COM) at touchdown to prevent excessive braking and allow a reduced ground contact time, with the latter being a well-documented characteristic of sprinting at higher speeds (9–11). The work of Mann and Herman (12) showed that greater top sprinting speed was associated with a reduced HTD, although this conclusion was based on an investigation of only three athletes. More recently, studies have shown that HTD does not appear to differ between athletes with different top sprinting speeds (13, 14) or to correlate with top sprinting speed (1). These conflicting findings make it unclear whether HTD is a kinematic variable coaches should target for sprinting.

**Figure 1.**
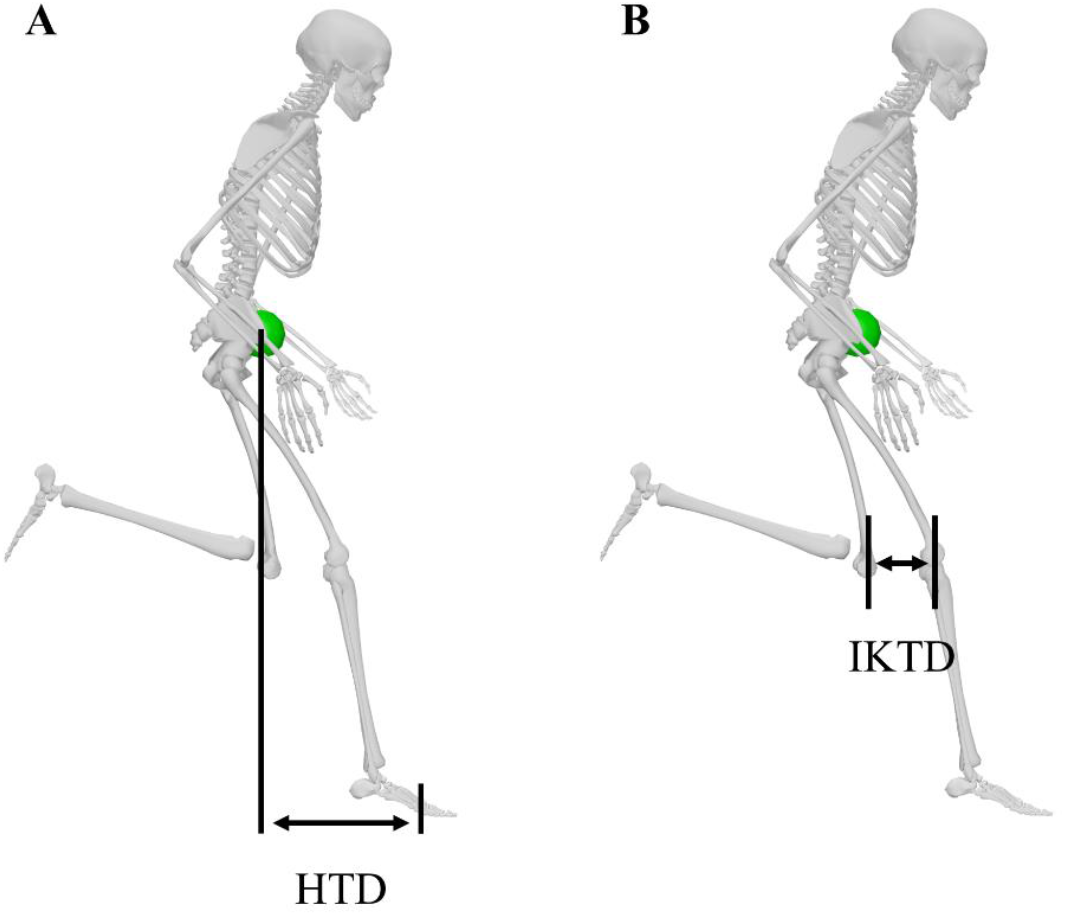
Commonly coached kinematic variables for sprinting. (A) Horizontal touchdown distance (HTD) is the horizontal distance between the whole-body center of mass (green sphere) and the foot at touchdown. (B) Inter-knee touchdown distance (IKTD) is the horizontal distance between the knees in the sagittal plane at touchdown.

Inter-knee touchdown distance (IKTD) (Figure 1B), an additional kinematic variable utilized in coaching, is part of the kinematics-based coaching framework for sprinting known as front-side mechanics (6–8, 15). The framework recommends a reduced IKTD, which should be achieved by ensuring that the hip of the swing limb is flexed at touchdown whilst the hip of stance limb is neutral at touchdown. The proposed benefit of reduced IKTD is that it enables the swing limb to contribute to vertical ground reaction force and impulse production in a manner analogous to how arm motions enhance jump height (1). Furthermore, a reduced IKTD, via greater hip flexion of the swing limb, is believed to permit the hip to achieve greater peak flexion, and thus a greater range of motion, leading to a greater extension velocity prior to ground contact (16). The combination of greater hip flexion-extension range of motion and hip extension velocity at touchdown have been shown to contribute to the hallmark rising-edge vertical ground reaction force production observed in athletes with greater top sprinting speeds (16, 17). Several studies have linked reduced IKTD to greater top sprinting speed (14, 18, 19), but Haugen et al. (1) found no correlation. Whilst the existing literature appears to show promise for IKTD as a key technique variable for sprinting, further work is warranted to explore whether IKTD has a substantial effect on speed.

To explore how HTD and IKTD affect top sprinting speed, studies have typically relied upon inter-athlete experimental study designs (1, 14, 18). However, this approach makes it difficult to isolate the influence of a single variable on speed due to the confounding contribution of varying physical attributes between athletes (20). While an intra-athlete experimental study design could help avoid these limitations, elite athletes and their coaches are reluctant to participate in interventions that alter biomechanics, given that kinematic changes may worsen performance or increase susceptibility to injury (21). Musculoskeletal simulation enables researchers to study kinematic modifications and their effect on top sprinting speed without the limitations of experimental approaches. For example, Rottier and Allen (19) developed a two-dimensional computer model to explore how the kinematics of the swing leg affect top sprinting speed, showing the promise of this approach

We used a three-dimensional predictive simulation approach to determine how commonly coached touchdown kinematics variables influence maximal running speed. In particular, we quantified changes in top speed as we enforced a range of HTD and IKTD values. We examined corresponding changes in ground reaction forces, spatiotemporal parameters, and lower-limb joint kinematics and kinetics to understand how the manipulations in the discrete touchdown kinematics variables influenced the top sprinting speed achieved in simulation.

## Methods

### Musculoskeletal Model

We used the three-dimensional, full-body musculoskeletal model described in Haralabidis et al. (22) (Figure 2), which is based on the model developed by Hamner et al. (23). A similar version of this model, together with a data-tracking optimal control framework, has been shown to reproduce joint kinematics, joint kinetics, and ground reaction forces with average root mean squared differences of 1°, 17.7 Nm, and 74.5 N, respectively, for maximal effort sprinting (22).

**Figure 2.**
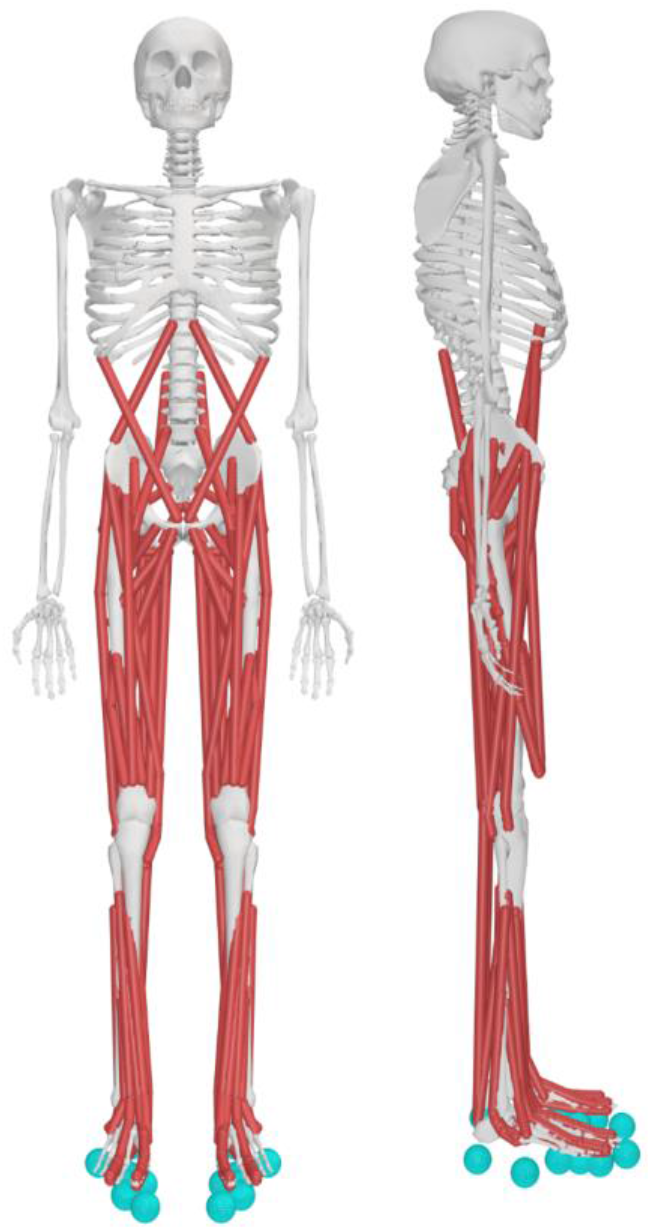
The musculoskeletal model described in Haralabidis et al. (22), adapted from Hamner et al. (23), used in this study. The skeleton has 20 rigid segments: a pelvis, trunk (torso plus head), right and left lower-limbs (thigh, shank, upper rearfoot, lower rearfoot, and forefoot) and upper-limbs (upper-arm, lateral forearm, medial forearm, and hand). The model’s 37 DOFs were: ×6 pelvis-to-ground, ×7 per lower-limb (×3 hip, ×1 knee, ×1 ankle, ×1 subtalar, and ×1 MTP), ×3 back and ×7 per upper-limb (×3 shoulder, ×2 elbow, and ×2 wrist). The MTUs and foot-ground contact elements are represented by the red lines and turquoise spheres, respectively.

The model represented the human body as a multibody system with 20 rigid segments and 37 degrees-of-freedom (DOFs). The lower-limb and trunk DOFs were actuated by 92 muscle-tendon units (MTUs) along with a torque actuator (40 Nm limit) at the metatarsophalangeal (MTP) DOF of each foot. We modeled each MTU as a three-element Hill-type model using the contraction and activation dynamics formulations of De Groote et al. (24) and De Groote et al. (25), respectively. The lengths, velocities, and moment arms of each MTU were described using differentiable and continuous polynomial functions (26). The upper-limb DOFs were actuated by 14 torque actuators (100 Nm limit) that included activation dynamics (27). A linear rotational spring at each of the MTP DOFs represented the passive properties of the forefoot and sprinting spikes, with the stiffness (65 Nm/rad) determined by summing prior literature values for sprinting spikes (28) and passive forefoot properties (29).

We modeled foot-ground contact by attaching four smooth Hunt-Crossley contact spheres (30) to each rearfoot segment and two contact spheres to each forefoot segment. We set the parameters of the contact spheres (location, uniform stiffness, and uniform damping) based on a prior investigation that involved calibrating the parameters specifically for sprinting (22). An additional contact sphere, included bilaterally at the heel of the rearfoot segment with the same parameters as the other contact spheres, prevented the heel from depressing beneath the ground. We applied an aerodynamic drag force to the pelvis segment at the whole-body center of mass using the approach of Samozino et al. (31).

We scaled the model linearly using the Scale Tool of OpenSim (version 3.3; Stanford University, CA, USA) (32) based on experimentally measured markers placed on anatomical landmarks to match the anthropometric and inertial characteristics of an international-level male sprinter (age: 24 years; height: 1.79 m; mass: 72.2 kg; 100 m Personal Best: 10.33 s; 200 m Personal Best: 20.27 s). The Hill-type model underpinning the force-generating capacity of each MTU is dependent on five parameters: pennation angle at optimum fiber length, tendon slack length, optimum fiber length, maximum shortening velocity and maximum isometric force. For this study, we used the pennation angle at optimum fiber length reported in the model of Hamner et al. (23), whilst we used the optimum fiber length and tendon slack length parameters obtained after scaling the model. We increased the maximum shortening velocity parameters from 10 to 12 optimal fiber lengths per second based upon an *in vivo* estimate (33). The maximum isometric force parameters were doubled with respect to those reported in the original model to more closely reflect the training status of the athlete the model was scaled to represent. This modification resulted in the model having a relative total individual lower-limb muscle volume of 119.5 cm^3^/kg by assuming a specific tension of 60 N/cm^2^, a tension value that is in accordance with the *in vivo* value reported for healthy young men (34); this volume falls within the *elite* category for male sprinters (126.3 cm^3^/kg) for comparable muscles (35).

The state variables of our model consisted of the generalized coordinates ***q***, generalized velocities ***v***, normalized MTU tendon forces 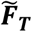, MTU activations ***a***, and upper-limb actuator activations ***a***_***UL***_. The control variables were the time derivative of the generalized velocities 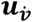, time derivative of normalized MTU tendon forces 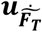, time derivative of MTU activations 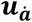, upper-limb actuator excitations ***e***_***UL***_ and reserve actuator activations ***u***_***R***_.

### Predictive Simulation Framework

We formulated predictive simulations of a symmetric sprinting step as optimal control problems and converted them to discrete nonlinear programming problems by using direction collocation, which features parameterization of both state and control variables. For each simulation, we discretized the step duration across 50 equally spaced mesh intervals, and within each mesh interval we further discretized at four *flipped* Legendre-Gauss-Radau (LGR) points (τ_*LGR*_ = [0.00, 0.155, 0.645, 1.00]) (36). We parameterized the state variables with third-order Lagrange polynomials at the LGR points, whilst we parameterized the control variables with second-order Lagrange polynomials at the two interior and terminal LGR points. We enforced MTU contractile and activation dynamics and multibody dynamics implicitly, by including 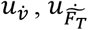,and 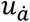 as control variables, and upper-limb activation dynamics explicitly. The implicit dynamics formulation resulted in a system of simple differential constraints in combination with algebraic path constraints (Table 1). We enforced all the model’s dynamics constraints at the two interior and terminal LGR points. In addition, we incorporated continuity constraints for the state variables between the ending and beginning of a pair of mesh intervals.

**Table 1.**
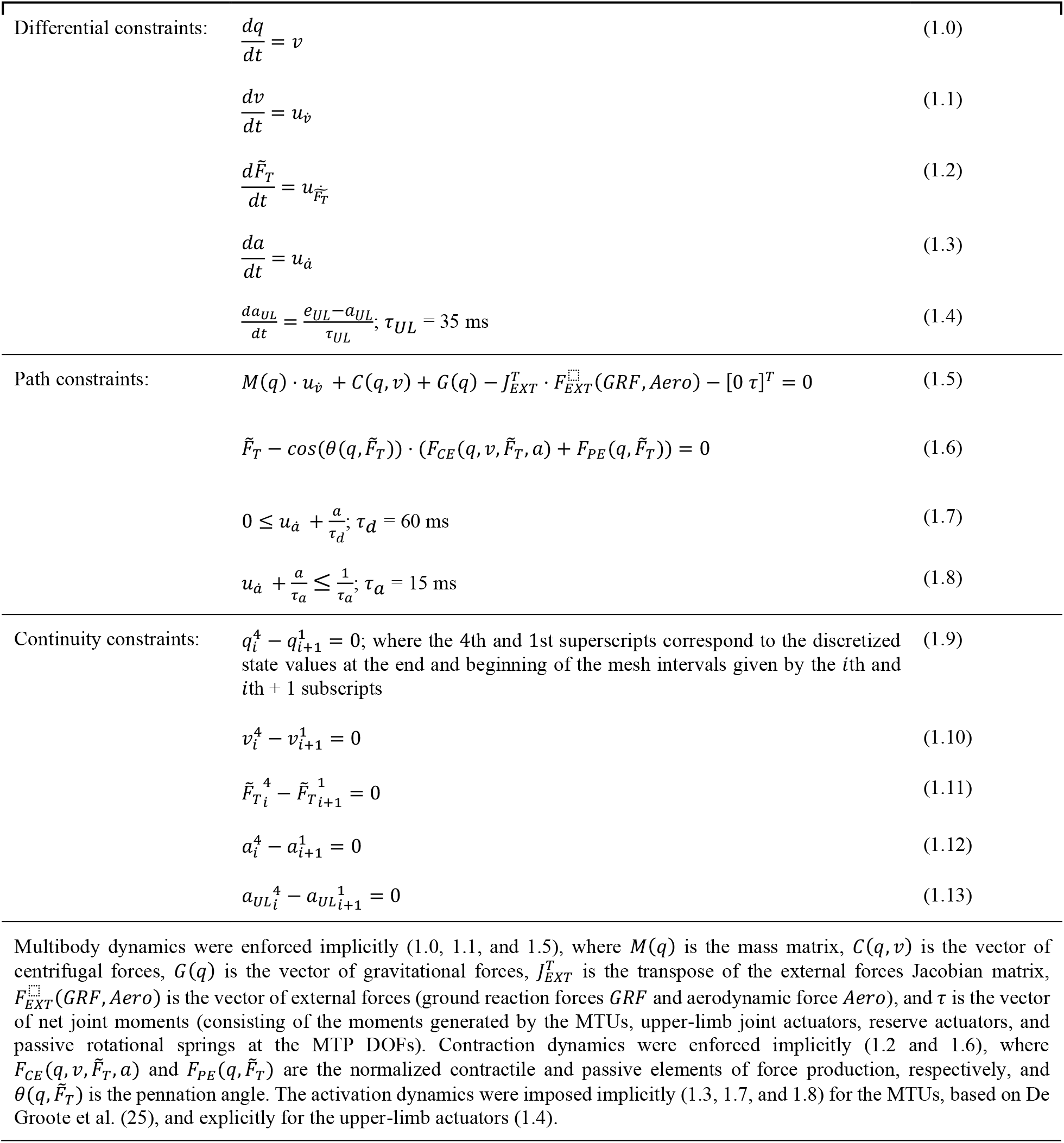
The musculoskeletal model’s dynamics constraints and state variable continuity constraints.

We incorporated additional task constraints across all predictive simulations. We set minimum distance inequality constraints, with a limit of 0.12 m, to prevent the shank and feet segments of opposing lower limbs from interpenetrating. We ensured the model’s configuration at touchdown reflected top-speed sprinting by imposing inequality constraints on the global pelvis and relative joint angles. We set the bounds for the initial configuration inequality constraints to ± 15° of previously determined global pelvis and relative joint angles at touchdown during top-speed sprinting for the athlete to whom the model was scaled (22).

We included an inequality constraint at the beginning of each simulation to ensure the vertical ground reaction force of the right foot was bounded between 20 and 40 N. This inequality constraint was included to ensure touchdown, which is commonly demarcated using a 20 N threshold (37, 38), occurred at the beginning of each simulation to enable exploring modifications to HTD and IKTD fairly across simulations. We enforced that the anterior-posterior pelvis translation was zero at the beginning of the step and could not exceed 2.47 m at the end of the step. The latter constraint was imposed to prevent solutions with an excessive step length, and the bound was set by multiplying the modeled athlete’s height with the maximum relative step length obtained in competition (39). Lastly, we simulated symmetric steps by enforcing that corresponding state (except anterior-posterior pelvis translation) and control variables at the beginning and end of a step matched.

We first performed a predictive simulation to determine the top speed of the model, hereafter referred to as the optimal simulation, and to identify baseline values for HTD and IKTD. We then performed a series of predictive simulations to explore how independently varying either HTD or IKTD affected speed.

These simulations were formulated as per the optimal simulation but also included an equality constraint to enforce either shortening or lengthening HTD or IKTD independently. We set the bounds for HTD and IKTD to ± 6 cm in 2 cm increments, leading to 12 predictive simulations in total (6 for HTD and 6 for IKTD). The bounds for varying HTD and IKTD were based on the standard deviation reported in prior literature (39, 40). We calculated a threshold to assess whether modifying HTD or IKTD resulted in meaningful changes to speed, and this was achieved by using the test-rest variation ratio of sprint speed (1.01) (41) together with the speed achieved in the optimal simulation (i.e., 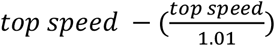.)

For each predictive simulation, we used an identical cost function *J* consisting of three terms:

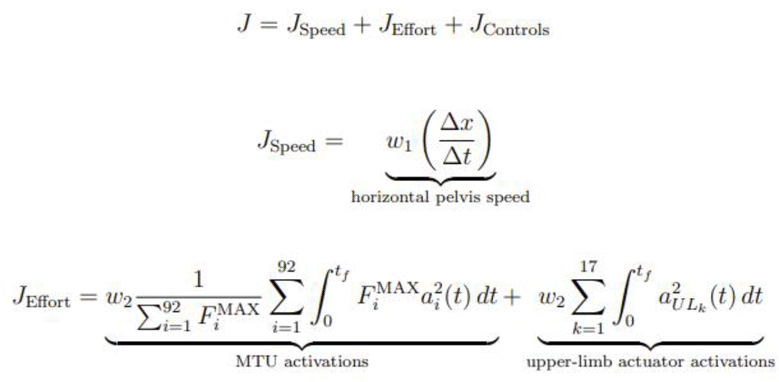

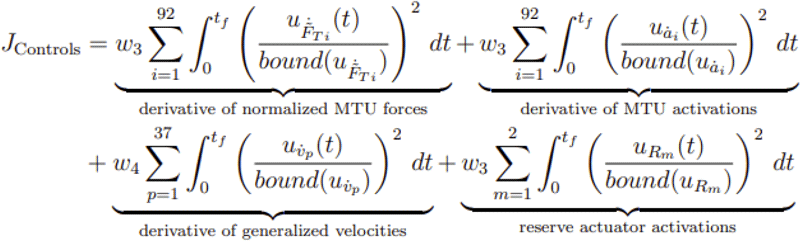

where *t*_*f*_ denotes the duration of a step and is a variable for each simulation, *w*_*i*_ are the weights of the cost function terms (*w* = [-10 0.1 0.01 0.05]), 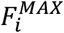 are the MTU maximal isometric force parameters, and *bound* represents the maximum absolute value of the lower and upper bounds of the parenthesized variable.

We formulated all simulations in MATLAB (2022b; MathWorks Inc., Natick, MA, USA) with CasADi (42), and solved them using IPOPT (43), an open-source nonlinear optimization solver, with an adaptive barrier parameter strategy and convergence tolerance of 10^−5^. We used versions of OpenSim and Simbody that have been modified to take advantage of CasADi’s algorithmic differentiation capabilities when evaluating the multibody dynamics (27). Details on the bounds and initial guess of the variables for all simulations can be found in Supplementary Material S1 and S2, respectively.

## Results

Our predictive simulation framework captured the salient features of elite, top-speed sprinting. In the optimal simulation (Figure 3 and Supplementary Video Material: https://github.com/nicos1993/Pred_Sim_Sprinting/tree/main/Videos), the model achieved a top speed of 11.85 m/s and a HTD and IKTD of 32.9 cm and 4.1 cm, respectively. The HTD and IKTD values in the optimal simulation are in the range of HTD (28 - 48 cm) and IKTD (0 - 7 cm) values reported in the literature (14, 15, 39). In addition, the kinematics, net joint moments, ground reaction force, and MTU activations of the optimal simulation reflected those of experimental data from top-speed sprinting (Supplementary Material S3). We set the threshold for a meaningful change in speed in response to varying either HTD or IKTD at 0.12 m/s based on the test-retest variation ratio of sprint speed and the speed achieved in the optimal simulation.

**Figure 3.**
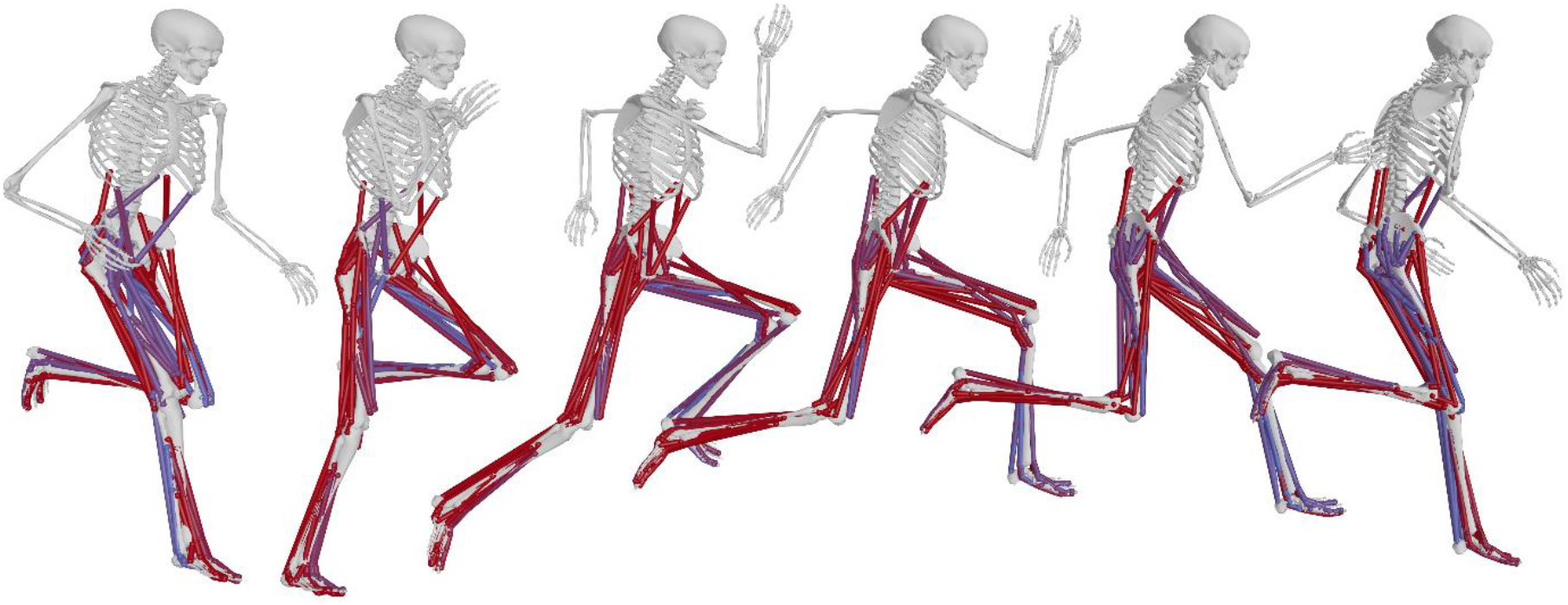
Sagittal plane visualization of the optimal simulation. For each MTU, the color represents simulated activation level from no activation (blue) to full activation (red).

We found meaningful reductions in speed when HTD was shortened by 2 cm or more and lengthened by 6 cm, with greater overall reductions in speed when shortening HTD (Figure 4A). We did not find meaningful reductions in speed when IKTD was modified (Figure 4B). For the remainder of the results, we focus on the effects of changing HTD by ± 6 cm; the effects of changing IKTD by ± 6 cm are included in Supplementary Material S4.

**Figure 4.**
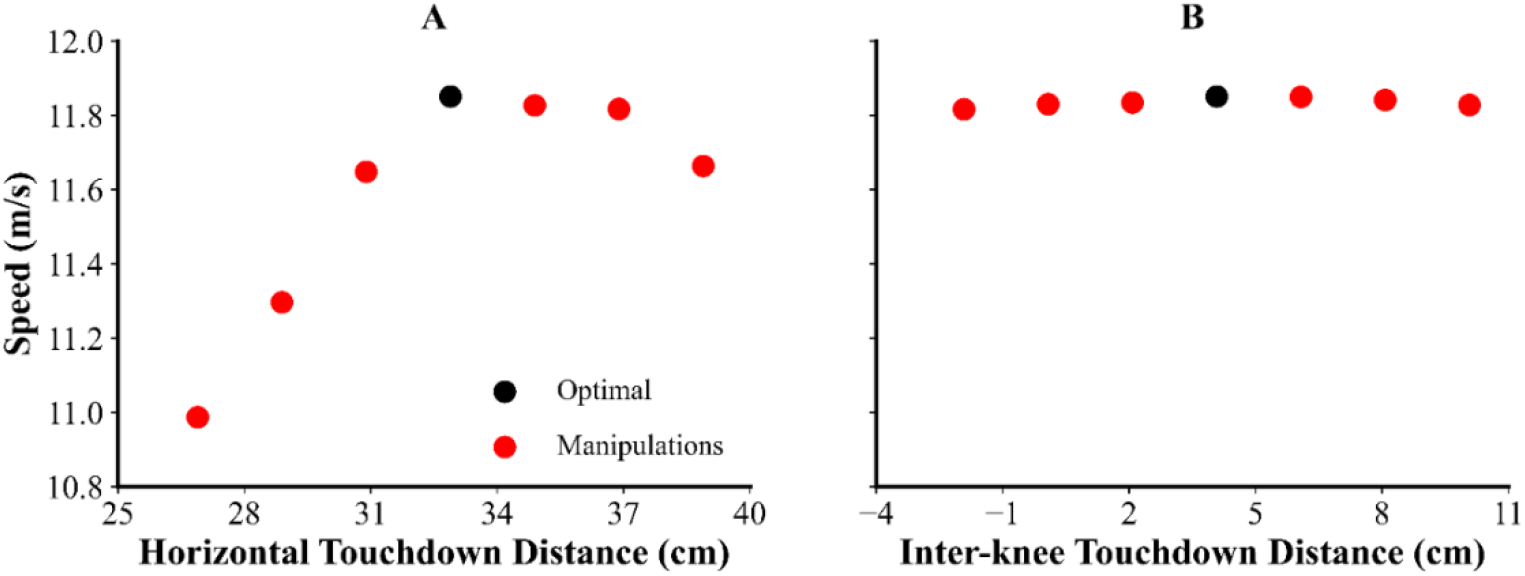
Horizontal touchdown distance (HTD) versus speed (A) and inter-knee touchdown distance (IKTD) versus speed (B). For both plots, the black circle is the optimal simulation, whilst the red circles are the simulations featuring HTD and IKTD manipulations.

### Shortening HTD

Shortening HTD by 6 cm reduced speed by 7.3%, and stance time by 7 ms (Table 2). Moreover, shortening HTD reduced net horizontal impulse and effective vertical impulse by 20.7% and 3.6%, respectively (Table 3 and Figure 5). We also observed differences in the hip joint kinematics of the stance and swing limbs (Figure S6), notably the stance hip was less flexed at touchdown and extended less during the stance phase. The temporal profiles of the joint moments were similar, although shortening HTD led to a greater peak stance ankle plantarflexion moment together with reduced stance knee flexion and hip extension moments (Figure S7). Interestingly, the net angular impulse of the stance ankle plantarflexion moment differed minimally (−15.0 vs. -15.1 Nms), whilst the net angular impulses of the stance knee flexion and hip extension moments were 0.5 Nms (−1.5 vs. -2.0 Nms) and 3.0 Nms (−0.8 vs. - 3.8 Nms) lower, respectively.

**Table 2.** Spatiotemporal variables from the optimal and largest horizontal touchdown distance (HTD) manipulation simulations.

| Simulation | Speed (m/s) | Step length (m) | Step frequency (Hz) | Contact time (ms) |
| --- | --- | --- | --- | --- |
| Optimal | 11.85 | 2.47 | 4.80 | 78 |
| - 6 cm | 10.99 | 2.16 | 5.08 | 71 |
| + 6 cm | 11.66 | 2.46 | 4.73 | 93 |

**Table 3.** COM velocity and ground reaction force variables from the optimal and largest horizontal touchdown distance (HTD) manipulation simulations.

| Simulation | Horizontal COM touchdown velocity (m/s) | Horizontal COM take-off velocity (m/s) | Vertical COM touchdown velocity (m/s) | Vertical COM take-off velocity (m/s) | Propulsive Impulse (Ns) | Net horizontal impulse (Ns) | Effective vertical impulse (Ns) |
| --- | --- | --- | --- | --- | --- | --- | --- |
| Optimal | 11.86 | 11.93 | -0.69 | 0.60 | 22.3 | 8.2 | 92.4 |
| - 6 cm | 10.99 | 11.05 | -0.68 | 0.56 | 18.7 | 6.5 | 89.1 |
| + 6 cm | 11.69 | 11.75 | -0.57 | 0.59 | 25.2 | 8.1 | 83.7 |

**Figure 5.**
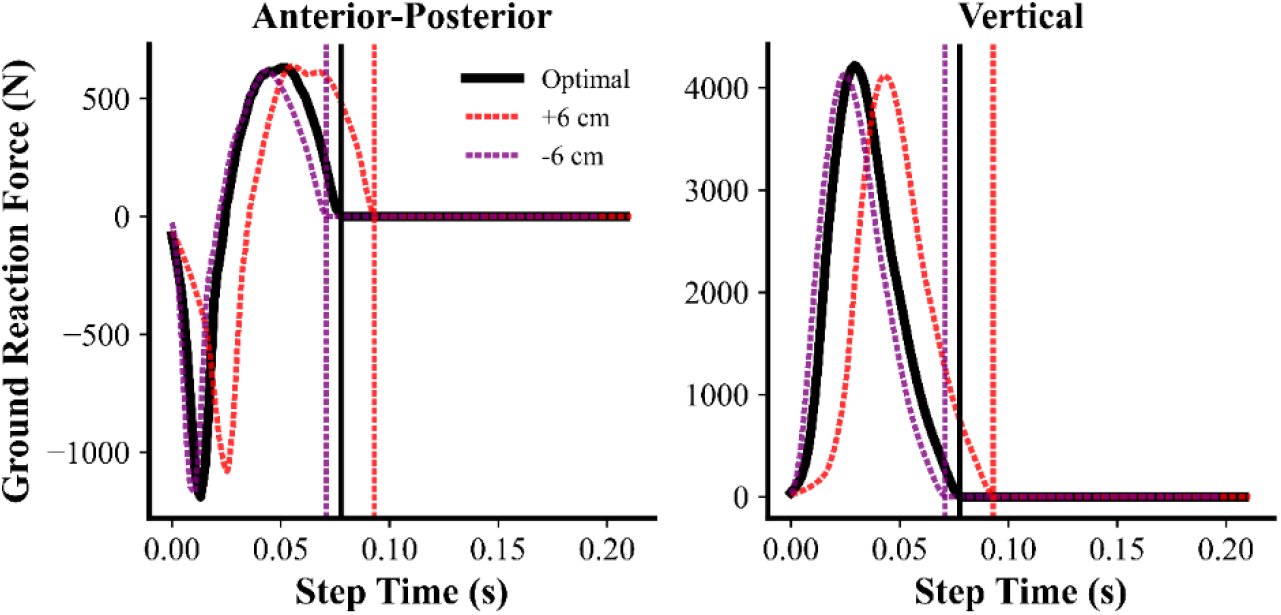
Ground reaction force time histories for the optimal simulation and the simulations with the largest horizontal touchdown distance (HTD) manipulation. Take-off for each simulation is indicated by the dotted vertical line. Data begin at touchdown of the right leg and end at touchdown of the left leg.

### Lengthening HTD

Lengthening HTD by 6 cm decreased speed by 1.6%, which was accompanied by a 15 ms increase in stance time (Table 2). Furthermore, lengthening HTD reduced the horizontal COM velocity at touchdown by 0.17 m/s (Table 3). We also observed differences in joint kinematics when HTD was lengthened (Figure S6), specifically for the stance ankle and knee and swing knee and hip. For the stance limb, lengthening HTD led to a lower peak ankle plantarflexion moment together with greater knee flexion and hip extension moments during the stance phase (Figure S7). The net angular impulse of the stance ankle plantarflexion moment was 0.5 Nms lower (−14.6 vs. -15.1 Nms) (Figure S7); however, the net angular impulses of the stance knee flexion and hip extension moments were 3.5 Nms (−5.5 vs. -2.0 Nms) and 6.5 Nms (−10.3 vs. -3.8 Nms) greater, respectively, in the simulation with lengthened HTD.

## Discussion

The simulated sprinter achieved a top speed comparable to elite male short distance track and field athletes (39), with kinematics, kinetics, and muscle activations that align with experimental results. We observed meaningful reductions in speed when we modified HTD, particularly when HTD was shortened. This suggests that there is an optimal HTD and that shorter is not necessarily better. When modifying IKTD, speed remained unchanged. These findings suggest that HTD is an important kinematic variable for top-speed sprinting, and warrants continued attention from coaches and athletes to optimize top speed. Conversely, IKTD may be less relevant, at least within the range we examined. Videos of the simulations, along with our code and corresponding data files can be accessed online at https://github.com/nicos1993/Pred_Sim_Sprinting.

Shortening HTD led to the largest decrease in speed (7.3%), and this coincided with lower ground reaction force and impulse production together with reduced ground contact time. Prior studies have identified that the ankle plantarflexor muscles contribute most to ground reaction force production during sub-maximal running (44) and maximal sprinting (45). But, interestingly, we found little change in the angular impulse of the net ankle plantarflexion moment when we shortened HTD, suggesting that a change in contribution of the ankle plantarflexor muscles was not the cause of reduced speed. We did, however, observe a reduced hip extension range of motion in the stance limb, which was accompanied by a smaller angular impulse of the net hip extension moment when we shortened HTD. Mann et al. (46) previously described a trade-off between ground contact time and ground reaction force production in terms of HTD, with the strength of the hip extensors (e.g., hamstrings and glutes) being a limiting factor given their role in pulling the COM over the foot during stance. Given that the model’s strength was identical for all the simulations performed, our findings suggest that shortening HTD reduced the capability of the hip extensor muscles to contribute to ground reaction force and impulse production, due to reduced net hip extension angular impulse, which led to the reduction in speed.

We also found a meaningful reduction in speed (1.6%) when we lengthened HTD. A longer HTD led to lower ground reaction force and impulse production along with increased ground contact time.

Furthermore, lengthening HTD resulted in greater braking impulse, a finding that coaches already recognize (7). This larger braking impulse led to an increased ground contact time in order to produce the necessary propulsive impulse to accelerate the COM and overcome air resistance. The angular impulse produced by the hip extension moment was greater when HTD was lengthened, perhaps to pull the COM forward, over the foot, through a larger distance. This coincided with the stance hip continuing to extend beyond take-off, which may have limited the role of the hip flexors late in stance and explains the lower hip flexion angular impulse generated. The reduced hip flexion angular impulse may have contributed to the reduction in sprinting speed given the important role that prior literature has placed on the hip flexion moment in the latter period of ground contact for reversing the rotation of the hip from extension to flexion (15, 47).

The reduced speed we observed when manipulating HTD differs from several prior studies (1, 13, 14). While the inter-athlete experimental designs in these past investigations may have masked the influence of changing HTD, our approach emulates an intra-athlete study design, allowing us to estimate how technique modifications affect performance on an individualized and isolated basis, which may be similar to how a coach would work with an athlete (8, 48). Mann and Sprague (49) suggested that an athlete with superior strength can overcome deficiencies in technique and vice versa. The approach used in the current study eliminates the confounding factor of strength, and permits focusing entirely on the technique aspects of sprinting performance.

Modifying IKTD had minimal impact on speed. Interestingly, we did observe an increase in vertical impulse production when IKTD was shortened, which supports the proposed mechanism underpinning shortened IKTD (1), although this did not translate to a faster top sprinting speed in our simulations. Our results highlight that there is sufficient redundancy in the neuromusculoskeletal system to maintain top-speed with a sub-optimal IKTD. In particular, the model achieved the targeted IKTD values by adjusting the swing-limb joint kinematics, whilst the kinematics of the stance limb remained largely unchanged. It is worthwhile noting that if an athlete is unable to achieve these changes in swing limb kinematics, altered IKTD could lead to reductions in top sprinting speed.

The minimal impact of IKTD on top sprinting speed we observed aligns with Haugen et al. (1), who found no relationship between IKTD (defined as the *inter-thigh angle*) and top sprinting speed based on an inter-athlete analysis featuring competitive sprinters. However, several prior studies have provided evidence in support of IKTD being a critical variable for top sprinting speed (14, 18, 19). Rottier and Allen (19) used a simulation approach to explore how the kinematics of the swing limb affect top sprinting speed and found a 0.9 m/s improvement in top sprinting speed when the kinematics of the swing limb were prescribed based on data from elite athletes. These prescribed kinematics had shortened IKTD. However, the conclusions from this study should be interpreted cautiously, as there was no guarantee that their model would have been able to execute the following step because they only simulated the stance phase. Our study did not have this limitation as we simulated an entire step and imposed symmetry.

While our study has provided valuable insights on top-speed sprinting, it is essential to acknowledge several limitations. Since our simulations relied on gradient-based optimization, the solutions we obtained may reflect a local rather than global optimum, and may be sensitive to the initial guess and variable scaling (50–52). To overcome the limitations of gradient-based optimization, we used the multistep approach of Ackermann and van den Bogert (53), first solving an optimization problem with several of the constraints relaxed, and then solving the problem again with the constraints incrementally tightened, using the solution from the prior optimization as an initial guess. We also initialized each optimization with multiple different guesses as in prior studies that used gradient-based optimization (51, 54, 55) and scaled each of the variables to be within the range of -1 to 1 to prevent poor numerical conditioning (50).

It should also be noted that we minimized MTU activations squared to resolve the MTU-force sharing problem in our cost function, which has been suggested to reflect MTU effort and energy consumption (53, 55); this may not be appropriate for top-speed sprinting. To evaluate the impact of this decision we compared a subset of the MTU activations determined from the optimal simulation to previously published electromyography (EMG) data for sprinting at speeds greater than 7 m/s (56). We observed generally good agreement in onset/offset timing (Figure S5). Our simulations did not activate the gastrocnemius and soleus in preparation of touchdown; however, this preparatory activation has not been consistently reported in prior studies of sprinting (45, 57). Lastly, despite using weighted MTU activations squared, the patterns of the major lower-limb joint moments in the simulations coincided with inverse dynamics analyses of sprinting (45, 58), which gives confidence in our findings.

While the focus of this study was top-speed sprinting, the ability of an athlete to accelerate is also vital to overall sprinting performance. In particular, our simulations did not include the steps preceding the attainment of top speed; instead we attempted to capture the effects of prior steps by loosely constraining the initial configuration of the model based on previous kinematics data for top-speed sprinting. However, in reality, an athlete’s top speed will be influenced by their preceding accelerative steps. Future work should, therefore, explore how the accelerative steps affect top sprinting speed, as well as the techniques associated with maximal accelerative ability.

In conclusion, we have shown that modifying HTD can lead to meaningful changes in top sprinting speed, whilst modifying IKTD did not. In the context of elite 100 m sprinting performance, top sprinting speeds of 11.85 and 11.73 m/s may translate to an 80 ms difference in overall 100 m time, based on the relationship between top sprinting speed and 100 m performance (2). Such differences can be the deciding factor between whether an athlete makes or misses the podium. Our results suggest that coaches should continue to place importance on HTD when working with athletes, meanwhile less effort should be devoted to improving IKTD. The predictive, high-speed sprinting framework that we developed could also be used to determine the impact of additional technique modifications on speed or injury risk.

## Acknowledgements

This work was supported by the Joe and Clara Tsai Foundation through the Wu Tsai Human Performance Alliance at Stanford University.

