## Supplementary Material for "Simulations reveal how touchdown kinematic variables affect top sprinting speed: implications for coaching"

***S1 Bounds***

We set the bounds of the generalized coordinates 𝒒, generalized velocities 𝒗, and derivative of generalized velocities $u_{\dot{v}}$ using maximal effort sprinting data of the athlete for whom the musculoskeletal model was scaled to (1). First, we performed inverse kinematics with OpenSim (version 3.3; Stanford University, CA, USA) (2) to determine the experimental generalized coordinates $q_{exp}(t)$. We then fit piecewise cubic splines to filtered $q_{exp}(t)$ (fourth-order low-pass Butterworth filter; 20 Hz cut-off frequency) to obtain approximated experimental generalized coordinates $\hat{q}(t)$. We then took the first and second derivatives of the polynomials with respect to time calculate the approximated generalized velocities $\hat{v}(t)$ and derivative of generalized velocities $\hat{u}_{\dot{v}}(t)$, respectively. In a penultimate step we determined the minimum ($\hat{q}_{min}$, $\hat{v}_{min}$ and $\hat{u}_{\dot{v}_{min}}$) and maximum ($\hat{q}_{max}$, $\hat{v}_{max}$ and $\hat{u}_{\dot{v}_{max}}$) values from $\hat{q}(t)$, $\hat{v}(t)$ and $\hat{u_{\dot{v}}}(t)$ across the entire trial. Lastly, we set the lower and upper bounds as follows:

$$\hat{q}_{min}-0.25\cdot\left| \hat{q}_{min} \right|\leq q\leq\hat{q}_{max}+0.25\cdot\left| \hat{q}_{max} \right|,$$

$$\hat{v}_{min}-0.25\cdot\left| \hat{v}_{min} \right|\leq v\leq\hat{v}_{max}+0.25\cdot\left| \hat{v}_{max} \right|,$$

$$\hat{u}_{\dot{v}_{min}}-0.25\cdot\left| \hat{u}_{\dot{v}_{min}} \right|\leq u_{\dot{v}}\leq\hat{u}_{\dot{v}_{max}}+0.25\cdot\left| \hat{u}_{\dot{v}_{max}} \right|,$$

In addition, we set the above lower and upper bounds to match bilaterally where necessary as we imposed symmetricity. We manually set the lower and upper bounds of anterior-posterior pelvis displacement $q_{pelvis-x}$ (units: meters):

$$0\leq q_{pelvis-x}\leq5$$

We set the upper bound of $q_{pelvis-x}$ to a value larger than permissible at an optimal solution, due to the constraint imposed on $q_{pelvis-x}$ (see Methods), to provide the optimization algorithm more freedom to explore at non-optimal intermediate iterations. We also manually set the lower and upper bounds of right and left upper-arm flexion-extension $q_{{UA}_{flex-ext}}$ and internal-external rotation $q_{{UA}_{int-ext}}$to prevent the arm segments from interpenetrating the torso and pelvis segments (which we observed in preliminary simulations):

$-75^{\circ}\leq q_{{UA}_{flex-ext}}\leq90^{\circ},$

$$-15^{\circ}\leq q_{{UA}_{int-ext}}\leq0^{\circ}$$

where the lower bounds corresponded to extension and external rotation, and the upper bounds corresponded to flexion and internal rotation. We set the lower and upper bounds of the step duration $t_{f}$ based on the experimental data (units: seconds):

$0.229-0.15\cdot0.229\leq t_{f}\leq0.229+0.15\cdot0.229$

We set the lower and upper bounds for MTU activations 𝒂, normalized MTU tendon forces $\tilde{F}_{T}$, time derivative of MTU activations $u_{\dot{a}}$, time derivative of normalized MTU tendon forces $u_{\dot{\tilde{F_{T}}}}$, upper-limb actuator activations $a_{UL}$ and excitations $e_{UL}$, and reserve actuator activations $u_{R}$as follows:

$$0 \leq a\leq1,$$

$$0 \leq\tilde{F}_{T}\leq5,$$

$$-\frac{\tau_{d}}{100} \leq u_{\dot{a}}\leq\frac{\tau_{a}}{100},$$

$$-1 \leq u_{\dot{\tilde{F_{T}}}}\leq1,$$

$$-1 \leq a_{UL}\leq1,$$

$$-1 \leq e_{UL}\leq1,$$

$$-1 \leq u_{R}\leq1$$

We set the bounds for the above variables, except $u_{R}$, based on the work of Falisse et al. (3).

***S2 Initial Guesses and Initialization***

We used a multistep approach, similar to Ackermann and van den Bogert (4), to determine the optimal simulation. We first performed an optimization in which the values of $q$, $v$, $u_{\dot{v}}$ and $t_{f}$ were initialized using the experimentally determined values (see above), the values of $e_{UL}$, $a_{UL}$ and $u_{R}$ were initialized as 0, $u_{\dot{\tilde{F_{T}}}}$ and $u_{\dot{a}}$as 0.01, and $\tilde{F}_{T}$ and 𝒂 as 0.5 and the symmetry constraints were imposed as inequalities with lower and upper bounds of 0.2 (all other constraints were enforced as described in the Methods). We then performed a further four optimizations in which we progressively tightened the bounds of the symmetry constraints (± 0.1, 0.05, 0.005, 0.0005), initializing all variable values using the prior optimal solution, after which a further optimization was performed with the symmetry constraints imposed as equalities. We then performed a final optimization, with symmetry equality constraints, to explore whether the optimal simulation could be improved upon using the variable values from the prior optimization as the initial guess. We defined the optimal simulation as the solution possessing the greatest horizontal speed from the latter two optimizations. We performed two sets of optimizations for each of the simulations exploring horizontal touchdown distance (HTD) and inter-knee touchdown distance (IKTD) modifications. For the first set of optimizations, we initialized all the variable values for each HTD and IKTD modification simulation based on the values determined from the optimal simulation. For the second set of optimizations, we initialized the variable values for each HTD and IKTD modification simulation based on the values determined from the first set of optimizations. We then used the solution possessing the greatest horizontal speed from the two sets of optimizations for each HTD and IKTD modification simulation for subsequent analysis.

***S3 Validation of Predictive Simulations***

We validated our predictive simulations by comparing the optimal simulation against previously published data for top-speed sprinting (1, 5), which is in accordance with the validation guidelines proposed by Hicks et al. (6). It is important to note that a direct comparison between simulated and experimental data was not possible (the speed achieved in the optimal simulation was 11.85 m/s whilst the speed for the experimental data ranged between 7 and 9.83 m/s), and this should be taken into consideration when assessing the simulated data.

The simulated vertical ground reaction force was found to have a slower rise time at the beginning of stance compared to the experimental data (Figure S1), and this difference can potentially be explained by Haralabidis et al. (1) using filtered experimental ground reaction force data, as opposed to raw data which is plotted, to calibrate the locations and parameters of the foot-ground contact model spheres. Additionally, the average and peak vertical ground reaction force differed by 0.41 (2.72 vs. 2.31 BW) and 2.05 (5.95 and 3.90 BW), and this can be attributed to the differences in speeds between the optimal simulation and experimental data, as prior experimental work (7) has identified an average vertical ground reaction force of approximately 2.5 BW for a speed similar to the speed achieved in the optimal simulation.

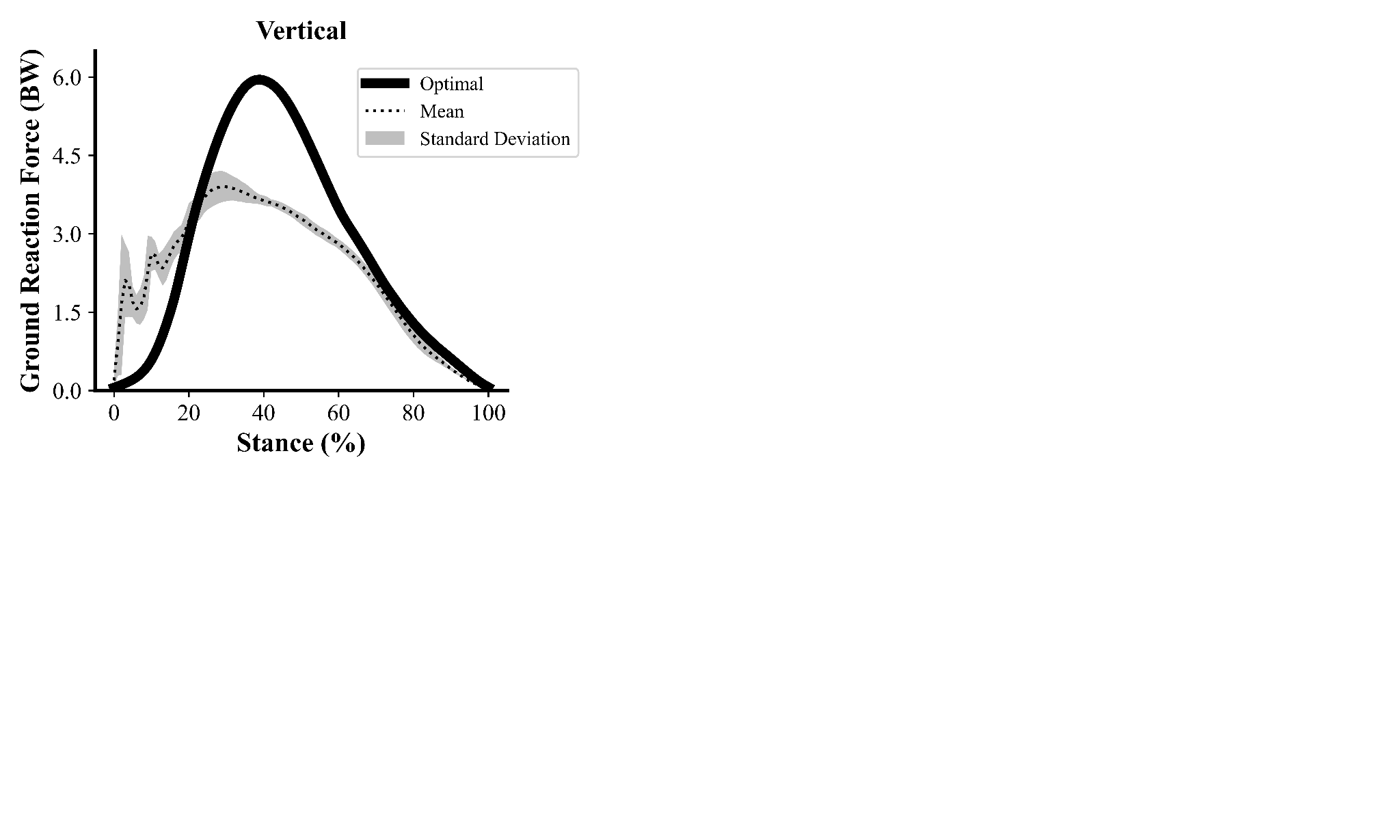

**Figure S1** Vertical ground reaction force during the stance phase for the optimal simulation (11.85 m/s) (solid black line) and experimental data (9.65 ± 0.25 m/s) from Haralabidis et al. (1) (dashed black line). Gray shaded areas are two standard deviations around the mean of the experimental data. Experimental data was not filtered.

The time histories of the major lower-limb joint and pelvis kinematics (Figure S2 and S3) for the optimal simulation were found to coincide with the experimental data for the most part. Notable differences included increased and decreased hip flexion and extension, respectively, for the lower-limb in stance at the beginning of the step for the optimal simulation, together with a more posteriorly tilted pelvis for the first half of the step.

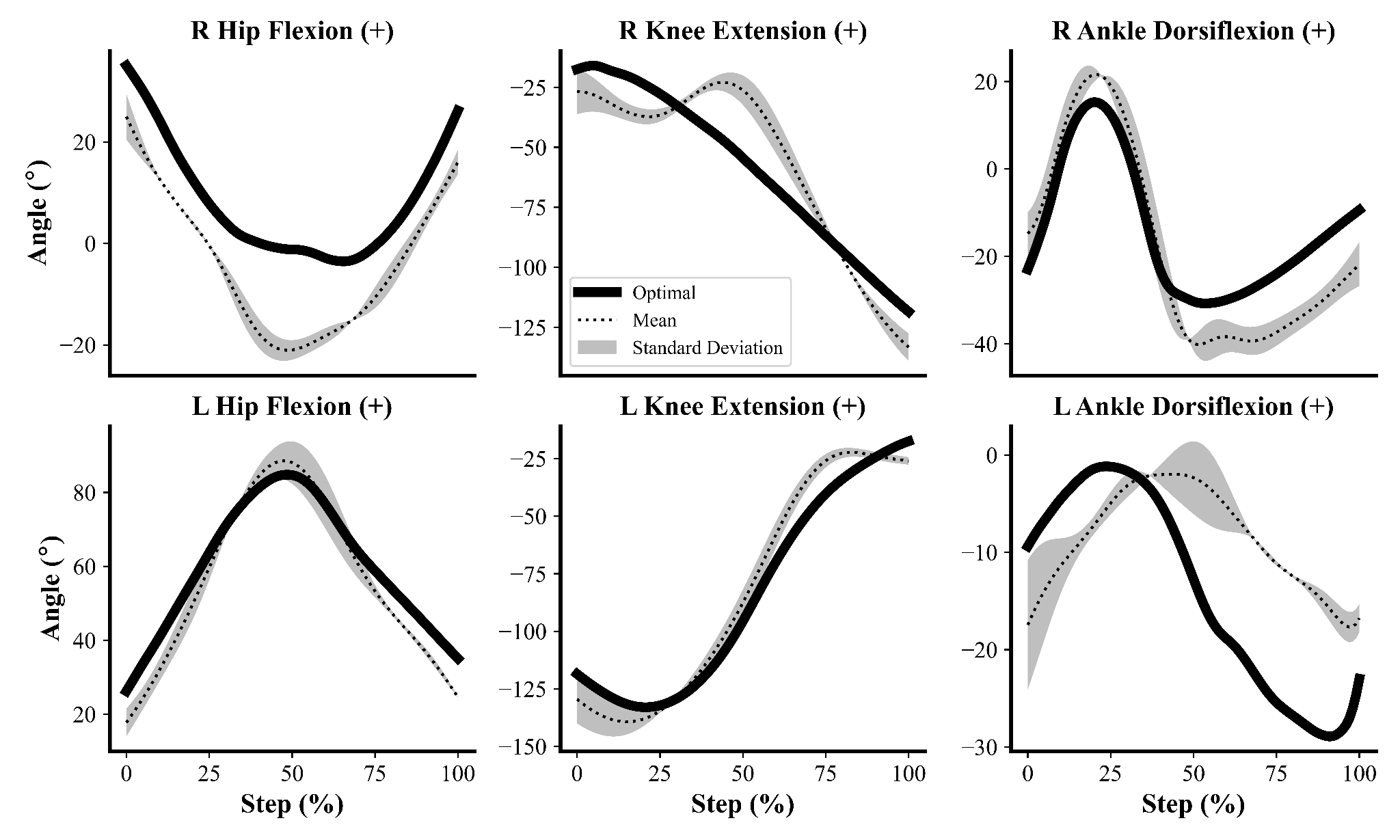

**Figure S2** Right (R) and left (L) major lower-limb joint kinematics during the step cycle for the optimal simulation (11.85 m/s) (solid black line) and experimental data (9.65 ± 0.25 m/s) from Haralabidis et al. (1) (dashed black line). Gray shaded areas are two standard deviations around the mean of the experimental data. Data begin at touchdown of the right leg and end at touchdown of the left leg.

The patterns of the net joint moments for hip flexion-extension and ankle dorsiflexion-plantarflexion of the stance limb in the optimal simulation coincided with those calculated from inverse dynamics analyses of the experimental data (Figure S4). Conversely, the net knee flexion-extension moment in the optimal simulation exhibited minimal extension throughout stance compared to the experimental data. It is worthwhile highlighting that this difference may be a consequence of the speed differences in the optimal simulation and the experimental data, as previous work has shown a decreased peak net knee extension moment as sprinting speed increases (8), leading to Bezodis et al. (9) speculating that there is a reduced reliance on the knee extension moment as speed increases.

For the swing limb, the pattern of the net hip flexion-extension moment in the optimal simulation followed the general trend from inverse dynamics, whilst the magnitude of the net ankle dorsiflexion-plantarflexion moment was similar between the optimal simulation and from inverse dynamics. Discrepancy in the pattern of the net knee flexion-extension moment for the swing limb between the optimal simulation and inverse dynamics was observed, although a previous study has identified a net knee extensor moment pattern for the swing limb whilst sprinting at 9.0 m/s (10) which therefore adds confidence to our results.

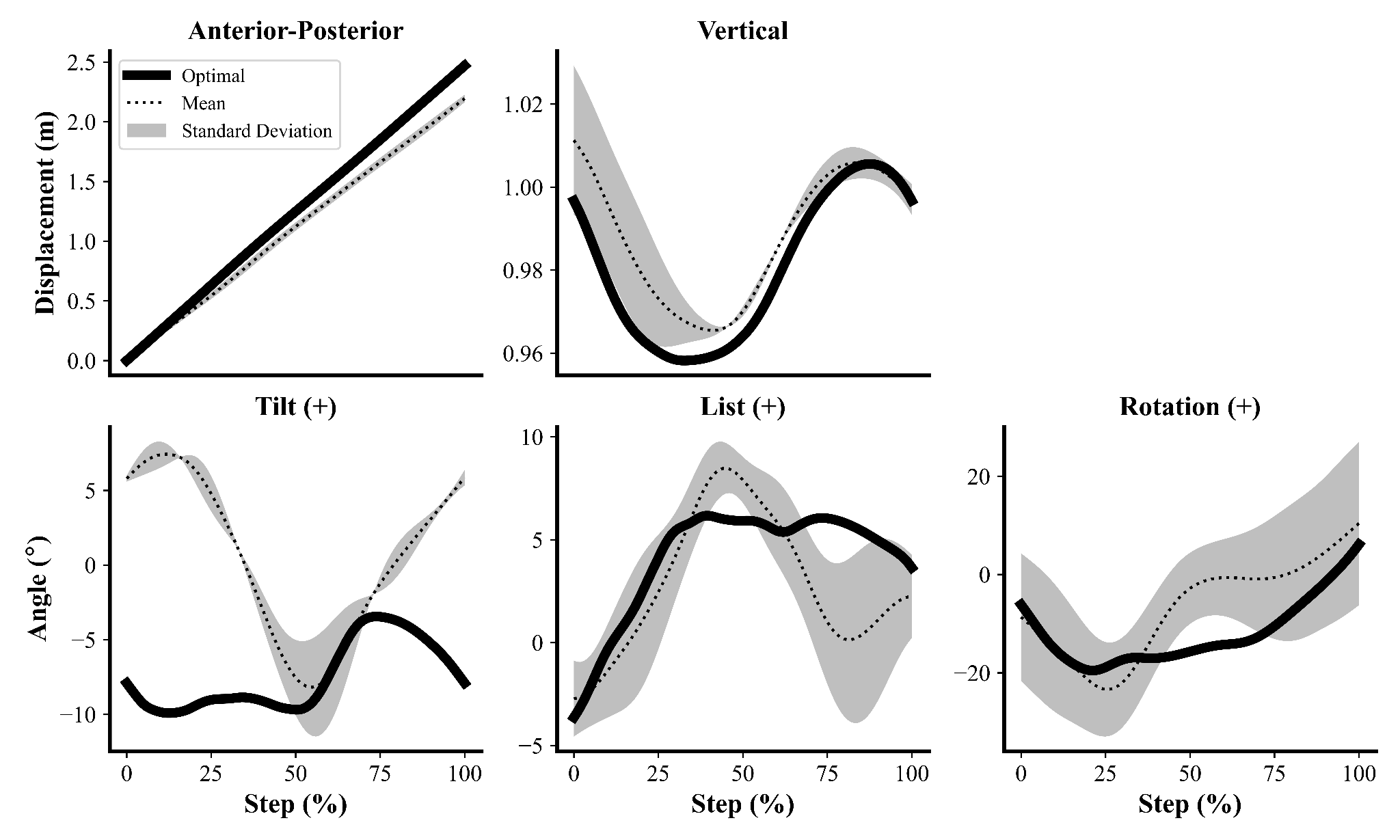

**Figure S3** Pelvis kinematics during the step cycle for the optimal simulation (11.85 m/s) (solid black line) and experimental data (9.65 ± 0.25 m/s) from Haralabidis et al. (1) (dashed black line). Gray shaded areas are two standard deviations around the mean of the experimental data. Data begin at touchdown of the right leg and end at touchdown of the left leg.

The periods of activation for a subset of MTUs throughout the stride cycle for the optimal simulation were found to be in agreement with experimental EMG data for the most part (Figure S5), although discrepancies were observed for the gastrocnemius, soleus and vastus lateralis in the late swing phase.

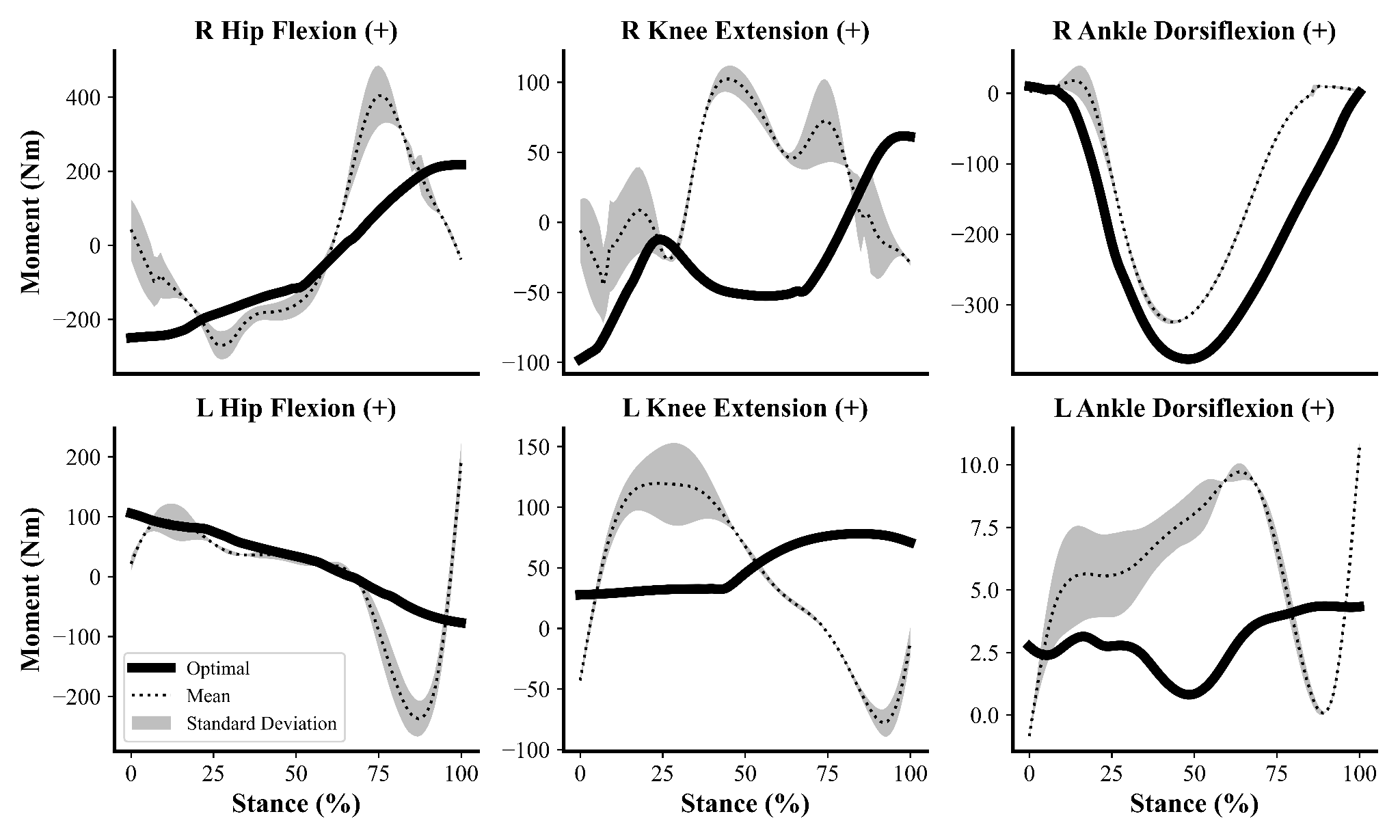

**Figure S4** Right (R) and left (L) major lower-limb net joint moments during the stance phase for the optimal simulation (11.85 m/s) (solid black line) and experimental data (9.65 ± 0.25 m/s) from Haralabidis et al. (1) (dashed black line). Gray shaded areas are two standard deviations around the mean of the experimental data. Data begin at touchdown of the right leg and end at take-off of the right leg.

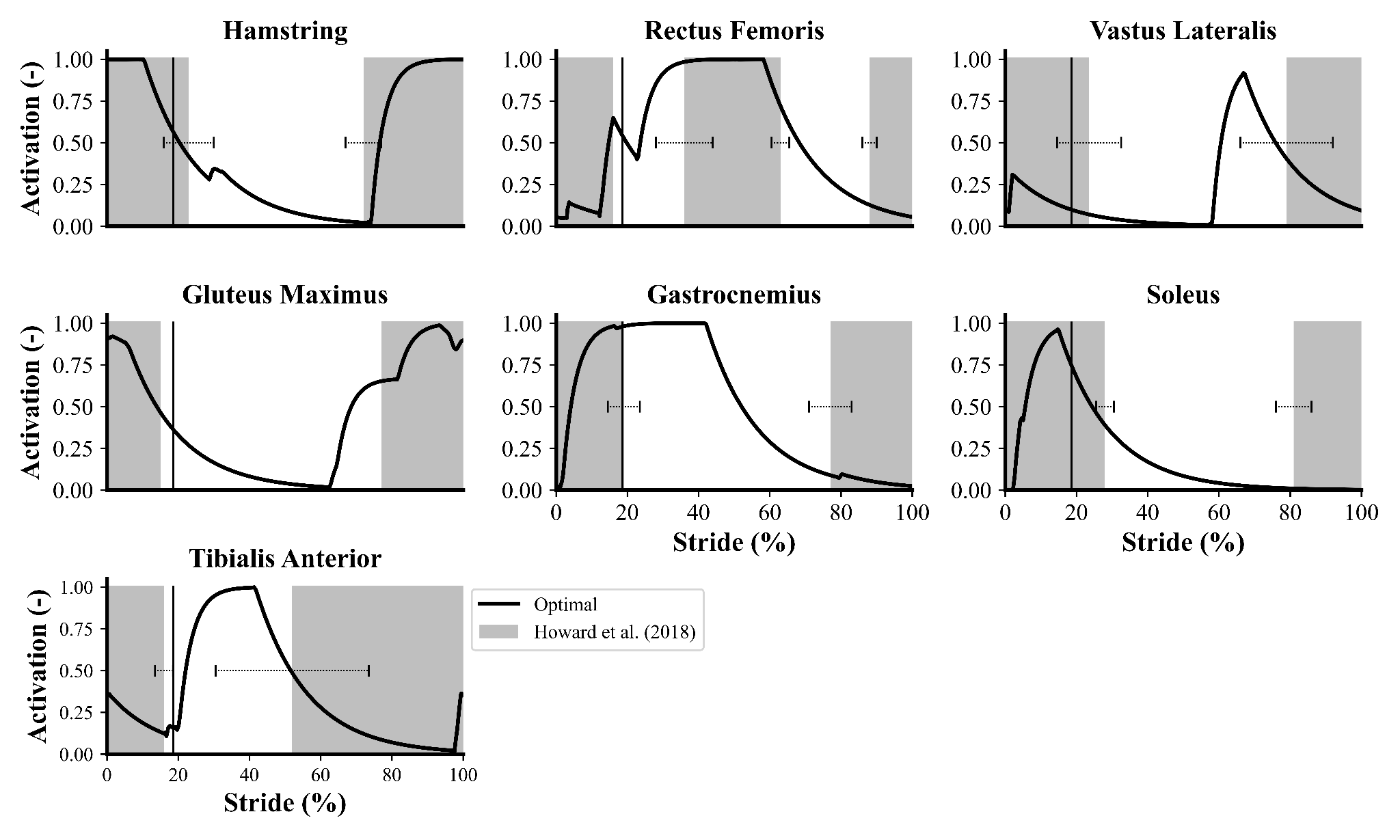

**Figure S5** Subset of MTU activations during the stride cycle for optimal simulation (11.85 m/s) (solid black line) (Hamstring: semimembranosus; Gastrocnemius: average of lateral and medial gastrocnemius), with take-off indicated by the thin vertical black line. The gray shaded regions and horizontal black tipped lines represent the mean and standard deviation (when available) periods of activation from experimental data (7.56 - 10.16 m/s) presented by Howard et al. (5).

***S4 Additional Results***

The time histories of the major lower-limb joint kinematics and net joint moments for the optimal simulation and those from the largest HTD manipulation simulations can be seen in Figures S6 and S7, respectively.

*Shortening HTD*

Shortening HTD by 6 cm resulted in a 0.87 m/s reduction in horizontal COM velocity at touchdown (Table 3). In addition, shortening HTD also resulted in a 0.04 m/s decrease in vertical COM velocity at take-off despite an almost equal vertical COM velocity at touchdown (Table 3).

*Lengthening HTD*

Lengthening HTD by 6 cm decreased net horizontal impulse by 0.1 Ns due to a greater braking impulse that had to be balanced by a greater propulsive impulse (Table 3 and Figure 5). We also observed a reduced effective vertical impulse, which culminated in a 0.01 m/s lower vertical COM velocity at take-off despite a less negative vertical COM velocity at touchdown (Table 3).

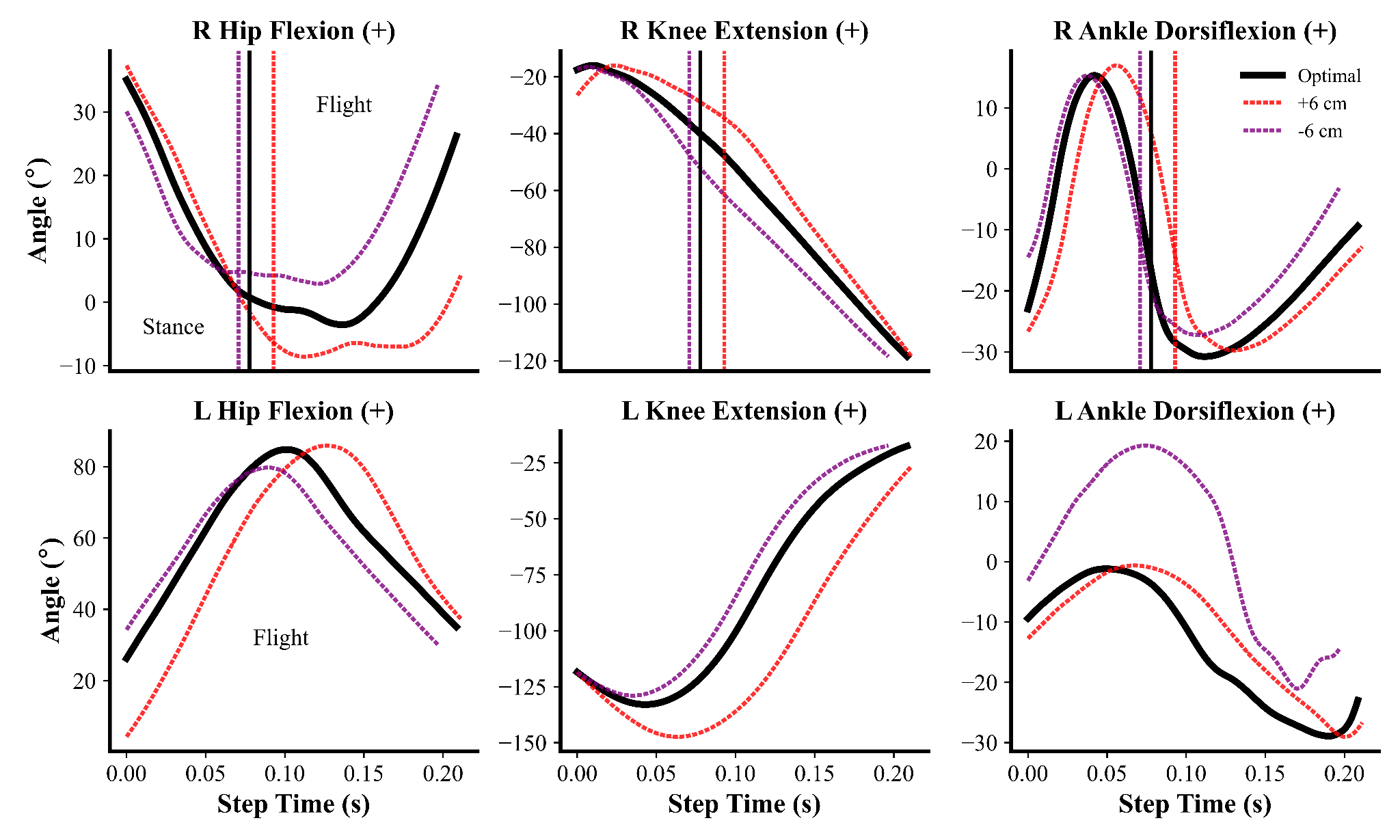

**Figure S6** Right (R) and left (L) major lower-limb joint kinematics time histories for the optimal simulation and for the simulations with the largest horizontal touchdown distance (HTD) manipulation. Data begin at touchdown of the right leg and end at touchdown of the left leg. Take-off for each simulation is indicated by the dotted vertical line, with stance and flight phases labeled for each lower-limb.

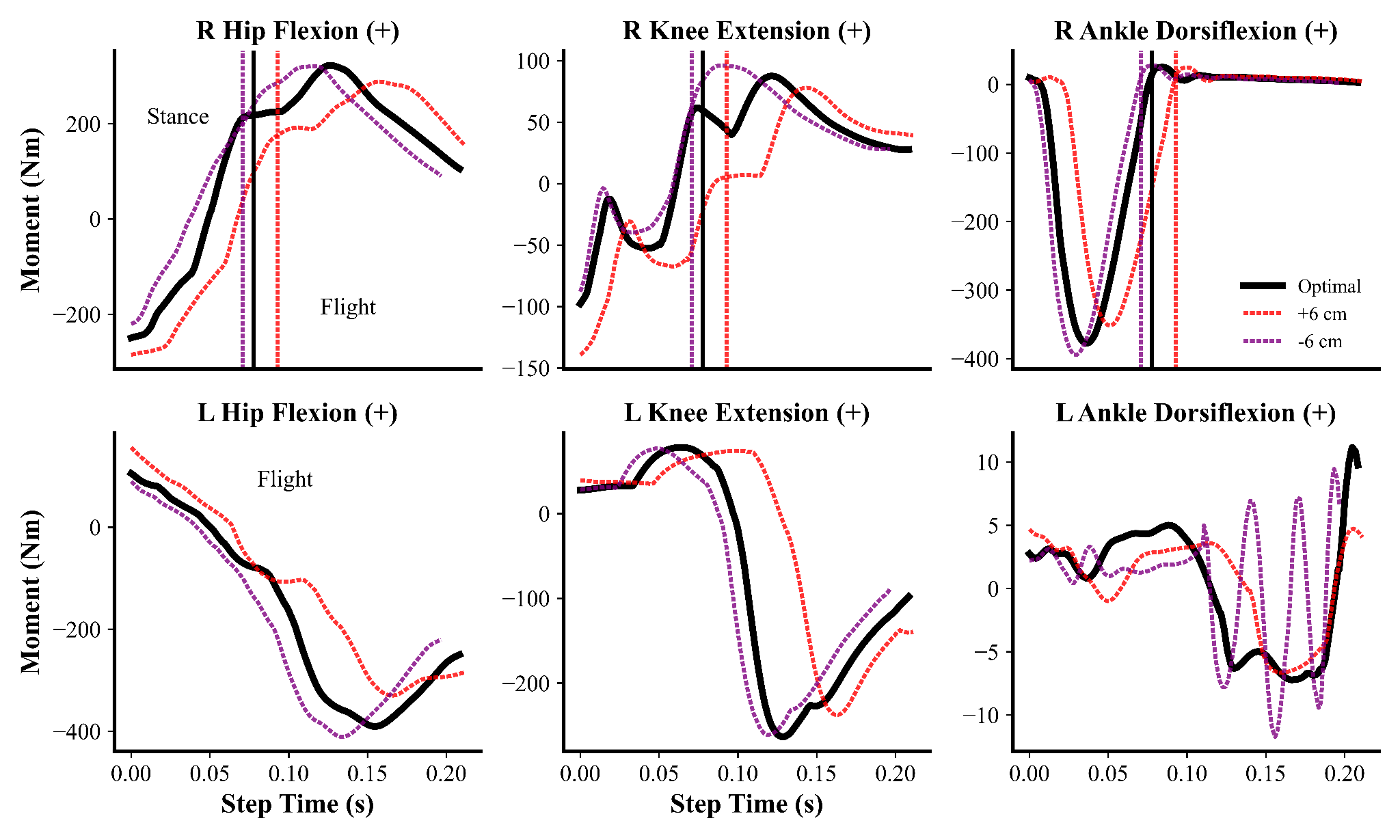

**Figure S7** Right (R) and left (L) major lower-limb joint moments time histories for the optimal simulation and those from the largest horizontal touchdown distance (HTD) manipulation simulations. Data begin at touchdown of the right leg and end at touchdown of the left leg. Take-off for each simulation is indicated by the dotted vertical line, with stance and flight phases labeled for each lower-limb.

*Shortening IKTD*

Shortening IKTD by 6 cm resulted in a 0.3% decrease in speed (11.82 vs. 11.85 m/s). There were minimal changes in spatiotemporal, COM velocity, and ground reaction force variables (Tables S1-S2 and Figure S8), with the exception of effective vertical impulse which increased by 2.2 N.s. Differences in the joint kinematics of the swing limb were observed when IKTD was shortened compared to the optimal solution (Figure S9), whilst for the stance limb they were visually indistinguishable. The swing hip was further flexed during stance whilst the knee remained more extended during the step. At the level of joint kinetics, the temporal profile of the stance and swing limb joint moments were similar when IKTD was shortened (Figure S10). However, for the stance limb, we observed a greater knee extension moment near take-off together with a reduced hip extension moment and greater hip flexion moment.

*Lengthening IKTD*

Lengthening IKTD by 6 cm led to a 0.2% reduction in speed (11.83 vs. 11.85 m/s). As for shortening IKTD, we observed minimal differences in spatiotemporal, COM velocity, and ground reaction force variables (Tables S1-S2 and Figure S8). Subtle differences were identified for the swing limb joint kinematics when IKTD was lengthened, with greater peak knee flexion and reduced hip flexion at touchdown (Figure S9). The stance limb joint kinematics were visually indistinguishable, except for the stance hip during the aerial phase which exhibited greater extension. The temporal profiles of the stance limb ankle dorsiflexion-plantarflexion and hip flexion-extension moments were similar, although we observed a reduced knee extension moment nearing the end of the stance (Figure S10).

**Table S1** Spatiotemporal variables from the optimal and largest inter-knee touchdown distance (IKTD) manipulation simulations.

| **Simulation** | **Speed (m/s)** | **Step length (m)** | **Step frequency (Hz)** | **Contact time (ms)** |
| --- | --- | --- | --- | --- |
| Optimal | 11.85 | 2.47 | 4.80 | 78 |
| - 6 cm | 11.82 | 2.47 | 4.78 | 75 |
| + 6 cm | 11.83 | 2.47 | 4.78 | 79 |

**Table S2** COM velocity and ground reaction force variables from the optimal and largest inter-knee touchdown distance (IKTD) manipulation simulations.

| **Simulation** | **Horizontal COM**  **touchdown**  **velocity (m/s)** | **Horizontal COM take-off velocity (m/s)** | **Vertical COM touchdown velocity (m/s)** | **Vertical COM take-off velocity (m/s)** | **Propulsive Impulse (Ns)** | **Net horizontal impulse (Ns)** | **Effective vertical impulse (Ns)** |
| --- | --- | --- | --- | --- | --- | --- | --- |
| Optimal | 11.86 | 11.93 | -0.69 | 0.60 | 22.3 | 8.2 | 92.4 |
| - 6 cm | 11.82 | 11.89 | -0.70 | 0.61 | 21.6 | 8.1 | 94.6 |
| + 6 cm | 11.84 | 11.91 | -0.68 | 0.59 | 22.7 | 8.2 | 91.6 |

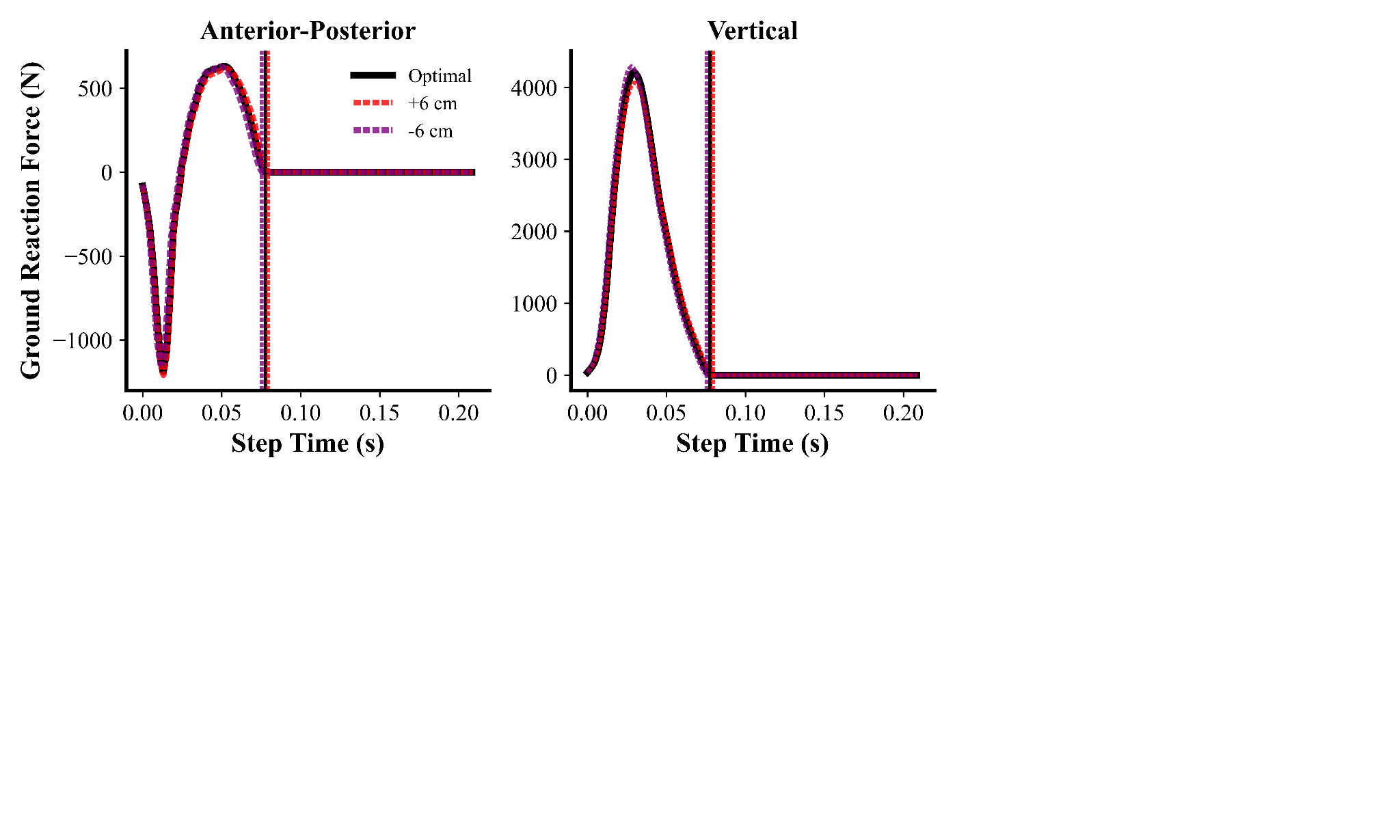

**Figure S8** Ground reaction force time histories for the optimal simulation and the simulations with the largest inter-knee touchdown distance (IKTD) manipulation. Take-off for each simulation is indicated by the dotted vertical line. Data begin at touchdown of the right leg and end at touchdown of the left leg.

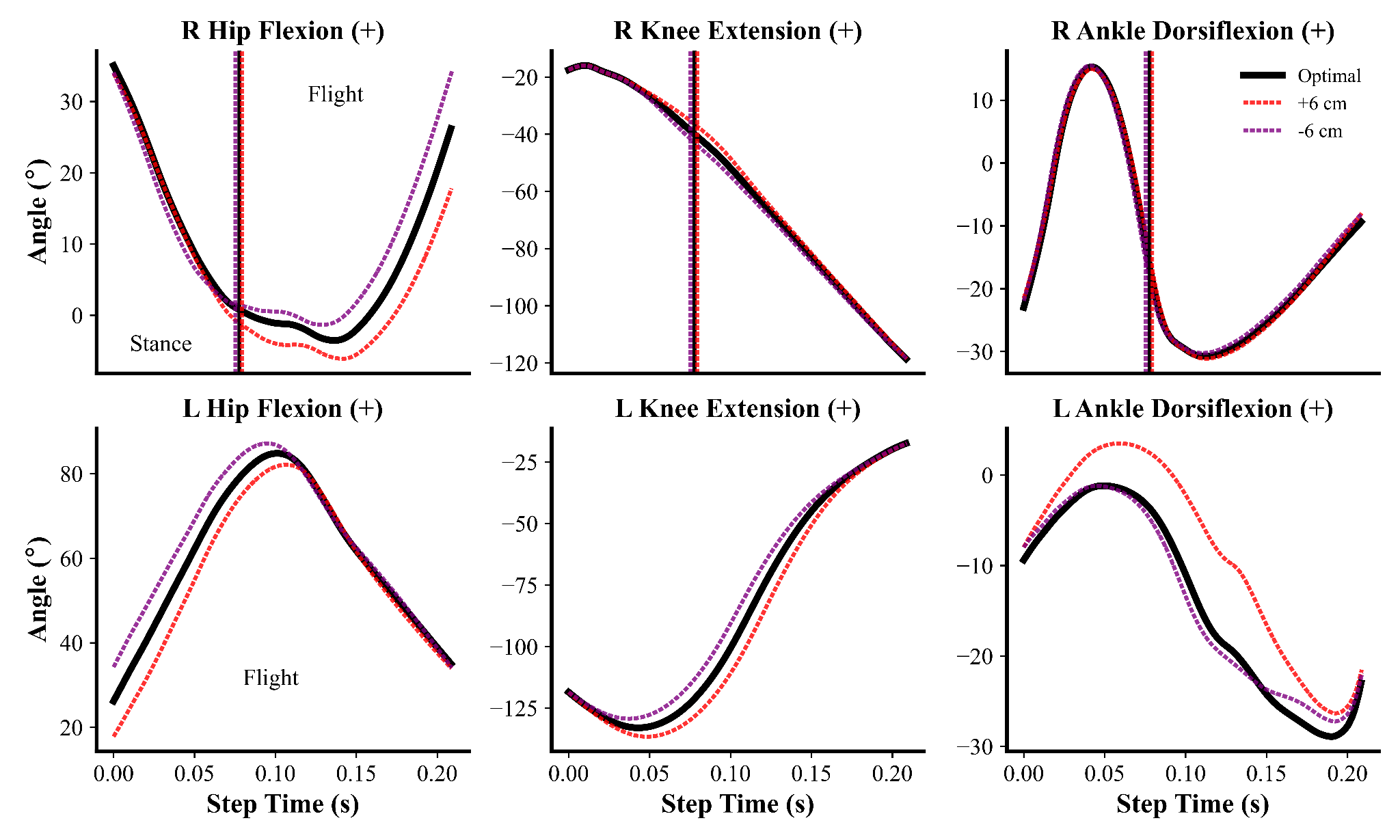

**Figure S9** Right (R) and left (L) major lower-limb joint kinematics time histories for the optimal simulation and those from the largest inter-knee touchdown distance (IKTD) manipulation simulations. Data begin at touchdown of the right leg and end at touchdown of the left leg. Take-off for each simulation is indicated by the dotted vertical line, with stance and flight phases labeled for each lower-limb.

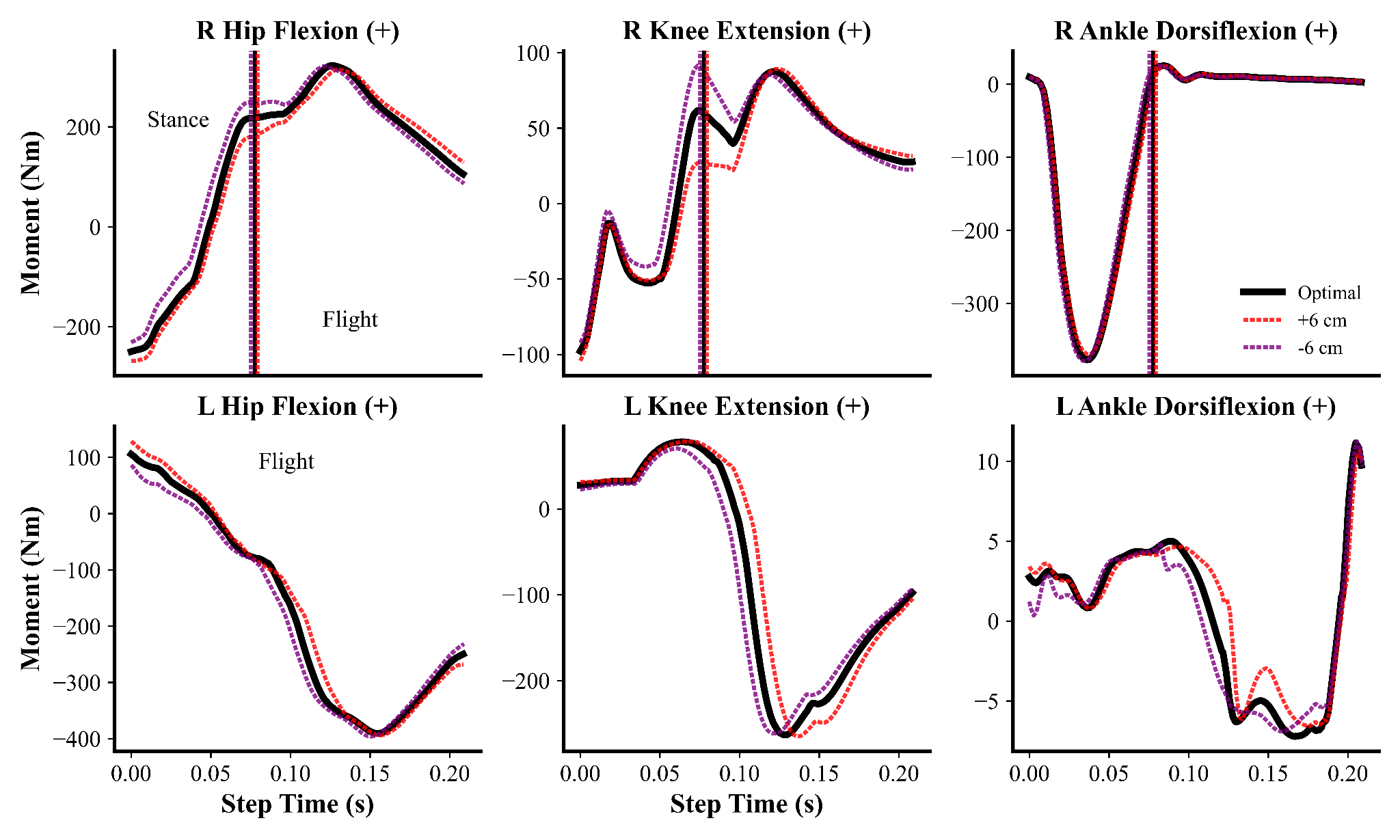

**Figure S10** Right (R) and left (L) major lower-limb joint moments time histories for the optimal simulation and those from the largest inter-knee touchdown (IKTD) manipulation simulations. Data begin at touchdown of the right leg and end at touchdown of the left leg. Take-off for each simulation is indicated by the dotted vertical line, with stance and flight phases labeled for each lower-limb.

**Supplementary Material References**
